# Poly(lactic-co-glycolic acid) immunomodulatory nanoparticles attenuate neuroinflammation and Alzheimer’s disease-related pathology in 5xFAD mice

**DOI:** 10.64898/2026.04.27.720934

**Authors:** Bryan Sanders, Matthew Korthauer, Kashish Singh Parihar, Igal Ifergan

## Abstract

Alzheimer’s disease is characterized by progressive cognitive decline, amyloid-β deposition, neuroinflammation, and neurodegeneration, yet effective and well-tolerated therapies remain limited. Because dysregulated myeloid responses are increasingly recognized as important drivers of disease progression, we investigated the therapeutic potential of poly(lactic-co-glycolic acid) immunomodulatory nanoparticles in the 5xFAD mouse model of amyloid-driven neurodegeneration. Poly(lactic-co-glycolic acid) immunomodulatory nanoparticles and fluorescently labeled particles displayed the expected size range and negative surface charge. After intraperitoneal administration, fluorescent particles were preferentially associated with myeloid cells in the blood, spleen, and brain, with greater uptake by brain myeloid populations in 5xFAD mice than in wild-type controls. Therapeutic treatment of 6.5-month-old 5xFAD mice, a stage at which behavioral abnormalities are already established, resulted in significant improvement in elevated plus maze behavior and a more modest improvement in Barnes maze performance. Flow cytometric analysis performed 9 weeks after the final treatment demonstrated persistent changes in brain immune composition, with the most prominent effects observed in P2RY12^+^ microglial populations, particularly the CD11c^+^ subset, and comparatively limited sustained effects in CD11b^+^P2RY12^-^ myeloid cells. These changes were accompanied by reduced expression of activation- and disease-associated markers and lower pro-inflammatory cytokine production within microglial populations. Histological analysis further showed reduced cortical amyloid plaque burden, decreased CD68 immunoreactivity, and reduced neurodegeneration in treated 5xFAD mice. Together, these findings show that systemically administered poly(lactic-co-glycolic acid) immunomodulatory nanoparticles produce durable behavioral, immunological, and pathological benefits in 5xFAD mice and support further investigation of this biodegradable myeloid-targeted platform as a therapeutic strategy for Alzheimer’s disease.

## 1. Introduction

Alzheimer’s disease (AD) is the most common neurodegenerative disease worldwide. Its clinical course typically begins with progressive impairment in learning and memory, followed by deficits in visuospatial function and language, worsening dementia, behavioral disturbances, and ultimately death, usually within 5–12 years after symptom onset [32, 50]. Two major pathological hallmarks define AD: the extracellular accumulation of amyloid beta (Aβ) peptides into plaques and the intracellular formation of neurofibrillary tau tangles. Aβ peptides are generated through sequential cleavage of amyloid precursor protein (APP), first by β-secretase (BACE1) and then by the γ-secretase complex, producing peptides of 38–42 amino acids in length. Among these, Aβ42 has the greatest propensity to aggregate and has been strongly implicated in neuronal injury [5, 32]. Neurofibrillary tangles arise from hyperphosphorylation of the microtubule-associated protein tau, leading to microtubule destabilization and neuronal dysfunction.

These two hallmarks have been the primary focus of AD research for several decades, and major efforts have been devoted to developing therapies that prevent or clear Aβ plaques and tau tangles. However, approaches that directly target these pathological features have so far yielded limited clinical benefit, and some therapies carry significant risks, including brain edema, microhemorrhage, neuronal toxicity, and amyloid-related imaging abnormalities (ARIA) [6, 42]. As the incidence of AD continues to rise with the aging population, there is an urgent need to develop safe and effective therapies that target additional drivers of disease progression.

There is growing consensus that neuroinflammation is a major contributor to AD pathogenesis. Epidemiological studies, transcriptomic analyses, and genome-wide association studies (GWAS) have implicated pro-inflammatory cytokines and dysregulation of genes associated with innate myeloid cells as important regulators of neurodegeneration in AD [4, 19, 30, 32, 51, 59]. In addition, multiple inflammatory signaling pathways linked to innate myeloid cells have been shown to be upregulated in AD, including activation of the NLRP3 inflammasome [3, 17]. Together, these findings support the idea that targeting inflammatory myeloid cells may represent a promising therapeutic strategy in AD.

Given the growing recognition of myeloid-driven neuroinflammation in AD, strategies that can selectively modulate immune cell function have gained increasing interest, with nanoparticles emerging as a particularly versatile approach. Their small size and tunable physicochemical properties make them useful for a wide range of applications, including diagnostics, antimicrobial therapy, tissue repair, and drug delivery [2, 20, 28, 46, 58]. Beyond these uses, nanoparticles can also exert immunomodulatory effects, as the inflammatory state of phagocytic immune cells is strongly influenced by uptake of exogenous material such as dead cells, lipids, microbes, and synthetic particles [11, 12]. Several nanoparticle platforms have been shown to suppress inflammatory programs in innate immune cells. For example, silver nanoparticles reduce inflammatory cytokine production in lipopolysaccharide-stimulated M1 macrophages [53], while cerium oxide nanoparticles decrease inducible nitric oxide synthase (iNOS) expression and reactive oxygen species (ROS) production in J774A.1 cells [18]. Although immunomodulatory nanoparticles (IMPs) can be generated from a variety of materials, highly biocompatible platforms are preferred in order to minimize toxicity and improve translational potential. For these reasons, we examined the therapeutic potential of poly(lactic-co-glycolic acid) immunomodulatory nanoparticles (PLGA-IMPs).

PLGA-IMPs are composed of lactic acid and glycolic acid monomers and offer several advantages over other nanoparticle platforms. PLGA is an FDA-approved biodegradable copolymer widely used in drug delivery systems, with multiple formulations already on the market [41]. Its degradation products can be metabolized through endogenous pathways including the tricarboxylic acid (TCA) cycle [34]. Consistent with this safety profile, PLGA-IMPs have recently been evaluated clinically for the treatment of celiac disease [26]. Recent studies indicate that PLGA-IMPs possess intrinsic immunomodulatory properties, particularly when formulated with carboxylated surface chemistry. These negatively charged nanoparticles are preferentially taken up by myeloid cells, while largely sparing resting immune populations and non-immune cells [22]. Following uptake, PLGA-IMPs reduce expression of pro-inflammatory markers and cytokines, likely by modulating key inflammatory signaling pathways [11, 29, 55].

Importantly, PLGA-IMPs have demonstrated therapeutic efficacy across multiple preclinical models of CNS inflammation and injury, including experimental autoimmune encephalomyelitis [11], spinal cord injury [24] and traumatic brain injury [45]. In these models, systemically administered PLGA-IMPs reduced infiltration of immune cells into the CNS, dampened neuroinflammation and reduced disease severity.

Recent work has demonstrated that native PLGA nanoparticles can attenuate AD-related pathology. In particular, intracerebroventricular delivery of native PLGA reduced amyloid burden and improved behavioral performance in young 5xFAD mice [1], while separate studies in primary neurons and iPSC-derived human neurons showed that native PLGA nanoparticles can regulate APP metabolism and protect against Aβ toxicity [54]. However, intracerebroventricular delivery is invasive and less readily translatable than systemic administration, underscoring the importance of determining whether PLGA-based particles can remain effective when delivered through a clinically more practical route. These findings support the therapeutic potential of native PLGA, but whether comparable benefit can be achieved through systemic administration at a later, behaviorally symptomatic stage of disease remains unclear. Here, we addressed this question by evaluating intraperitoneal PLGA-IMP administration in 6.5-month-old 5xFAD mice with established behavioral abnormalities, with particular focus on behavioral outcomes, myeloid-cell targeting, neuroinflammation, and downstream pathological changes.

## 2. Materials and Methods

### 2.1 Mice

5xFAD mice were generated by crossing hemizygous 5xFAD mice (MMRRC stock no. 34848) with C57BL/6J wild-type (WT) mice (The Jackson Laboratory stock no. 000664) to produce 5xFAD transgenic mice and non-transgenic littermate controls. Genotyping was performed by Transnetyx using ear tissue samples and PCR-based detection of the human APP Swedish (K670N/M671L) and PSEN1 (M146L/L286V) transgenes, as previously described [38]. Mice were bred and maintained under specific pathogen-free conditions in the University of Cincinnati Laboratory Animal Medical Services (LAMS) facility. All procedures were performed in accordance with institutional guidelines and were approved by the Institutional Animal Care and Use Committee (IACUC). Both male and female mice were used unless otherwise specified.

### 2.2 PLGA-IMP and FITC-PLGA-IMP synthesis

PLGA-IMPs were prepared using a single-emulsion solvent evaporation method adapted from previously described protocols [21, 37, 48]. Briefly, 400 mg of acid-terminated 50:50 poly(D,L-lactide-co-glycolide) (PLGA; Polysciences, cat# 26269; inherent viscosity 0.16–0.24 dL/g) was dissolved in 2 mL dichloromethane. The polymer solution was emulsified in 10 mL of 1% (w/v) Poly(ethylene-alt-maleic anhydride) (PEMA; Sigma) by probe sonication on ice for 30 s at 40% amplitude. Immediately after sonication, the emulsion was transferred into 200 mL of 0.5% (w/v) PEMA under stirring and left overnight to allow complete solvent evaporation. Nanoparticles were collected by centrifugation at 7,000 x g for 15 min and washed three times in 0.1 M sodium carbonate-sodium bicarbonate buffer (pH 9.6). Particles were then frozen at −80°C, lyophilized, and stored at −20°C until use. The acid-terminated 50:50 PLGA used in these studies is specified by the manufacturer as having 47–53 mol% glycolide, 47–53 mol% DL-lactide, and an acid number > 6 mg KOH/g.

FITC-PLGA-IMPs were prepared using the same method, except that the polymer phase consisted of 300 mg unlabeled PLGA mixed with 100 mg FITC-conjugated PLGA (NanoSoft Polymers). All subsequent emulsification, washing, freezing, and lyophilization steps were performed as described above.

Before in vivo use, lyophilized PLGA-IMPs and FITC-PLGA-IMPs were rehydrated in sterile PBS, washed three times with sterile PBS, and passed through a 40 μm filter to reduce aggregation. Particles were vortexed immediately before injection. Particle size distribution and Zeta-potential were determined by dynamic light scattering using a Zetasizer Nano ZS (Malvern Instruments). For these measurements, reconstituted nanoparticles were suspended in PBS, diluted in nanopore water at a concentration of 1 mg/mL, and analyzed immediately. Batches selected for *in vivo* studies displayed a dominant primary peak within the expected size range, with only minimal contribution from secondary peaks.

### 2.3 In vivo treatment

For therapeutic studies, 6.5-month-old 5xFAD mice received intraperitoneal (i.p.) injections of either PLGA-IMPs (2.5 mg in 200 μL PBS) or vehicle control (200 μL PBS) three times per week for 4 weeks. For particle-uptake studies, WT and 5xFAD mice received a single i.p. injection of FITC-PLGA-IMPs (2.5 mg in 200 μL PBS), and blood, spleen, and brain tissues were harvested 6 h later for cell isolation and analysis. PLGA-IMPs and FITC-PLGA-IMPs were vortexed immediately before each injection to minimize particle aggregation. Non-transgenic littermate controls (WT) used in therapeutic studies did not receive injections and served as untreated controls.

### 2.4 Behavioral testing

Behavioral tests were performed at 3 and 9 weeks after the final treatment injection.

#### Elevated plus maze

Mice were acclimated to the behavioral testing room for at least 30 min before testing. At the start of the assay, each mouse was placed in the center of the maze facing an open arm and allowed to explore freely for 5 min. The maze was cleaned between trials using 10% Decon solution, rinsed with water, and dried. Total distance traveled and time spent in the open arms were recorded and analyzed using AnyMaze software (version 7.48). Mice that fell from the maze at any point during testing were excluded from analysis.

#### Barnes maze

Barnes maze testing was performed over 3 consecutive days. On day 1, mice were habituated to the maze by being placed in the center of the platform for 60 s and then gently guided to the escape hole, where they remained in the escape chamber for an additional 60 s. On day 2, mice underwent 4 training sessions under aversive conditions (bright light and fan) and were given up to 3 min per session to locate the escape chamber. Mice that did not locate the escape hole within 3 min were guided to it, remained in the chamber for 60 s before being returned to their cages, and were assigned a latency of 180 s. Mice that successfully located the escape hole were allowed to remain in the chamber for 30 s before being returned to their cages. Training sessions were separated by 20 min between sessions 1 and 2 and between sessions 3 and 4, with a 4 h interval between sessions 2 and 3. On day 3, mice were placed in the center of the maze and allowed to explore for up to 90 s, and latency to locate the escape hole was recorded. For the 9-week assessment, the location of the escape hole was moved to the opposite quadrant relative to the 3-week test. Mice were acclimated to the behavioral testing room for at least 30 min before testing each day. Maze activity was recorded and analyzed using AnyMaze software (version 7.48). The maze and escape chamber were cleaned between trials with 20% ethanol, rinsed with DI water, and dried. Mice that fell from the maze during testing were excluded from analysis.

### 2.5 Tissue collection and leukocyte isolation Brain leukocyte isolation

For analysis of the therapeutic effects of PLGA-IMPs, brains were collected at the 9-week time point after the final treatment injection, following completion of behavioral testing. Mice were euthanized with CO₂ and transcardially perfused with 30 mL ice-cold PBS to remove circulating blood cells. Brains were harvested, mechanically dissociated, and enzymatically digested in a 1:1 mixture of DMEM and PBS containing collagenase (1 mg/mL; Worthington) and DNase I (0.1 mg/mL; Sigma-Aldrich) for 1 h at 37°C. Cell suspensions were passed through 100 μm strainers, and mononuclear cells were isolated by density centrifugation using 20% BSA to remove myelin and debris. Cells were then washed and resuspended in complete RPMI-1640 containing 10% FBS, 2 mM L-glutamine, and 100 U/mL penicillin-streptomycin for downstream flow cytometric analysis.

#### Splenocyte isolation

Spleens were harvested from perfused mice and mechanically dissociated through 100 μm strainers. Red blood cells were lysed using 0.83% ammonium chloride. Cells were then washed and resuspended in PBS for flow cytometric analysis.

#### Peripheral blood leukocyte isolation

Whole blood was collected at the time of euthanasia, and red blood cells were lysed using 0.83% ammonium chloride. The remaining leukocytes were washed and resuspended in PBS for flow cytometric analysis.

### 2.6 Flow cytometry staining and analysis

Single-cell suspensions isolated from brain, spleen, and blood were incubated with anti-mouse CD16/32 Fc block prior to staining. Cells were then stained with a fixable LIVE/DEAD viability dye (Invitrogen) for 20 min at room temperature in the dark, followed by surface staining for 20 min at 4°C. Surface markers included antibodies against CD45, CD11b, P2RY12, CD11c, CD80, CD86, and TREM2. For experiments assessing intracellular cytokine expression, cells were subsequently fixed and permeabilized using a fixation/permeabilization kit (Thermo Fisher Scientific) and stained for IL-1β, TNF-α, IL-10 and IL-12p40. Antibodies were obtained from BioLegend, BD Biosciences, Thermo Fisher Scientific (including Invitrogen and eBioscience), and R&D Systems.

For FITC-PLGA-IMP uptake experiments, brain myeloid populations were analyzed based on CD45 and CD11b expression and are reported as CD45^int^CD11b^+^ and CD45^hi^CD11b^+^ cells (**Fig. S1**). For treatment-response experiments, brain CD45^+^CD11b^+^ populations were further classified based on P2RY12 and CD11c expression and are referred to here as P2RY12^+^ microglia, P2RY12^+^CD11c^+^ microglia, and CD11b^+^P2RY12^-^ myeloid cells (**Fig. S2**).

When intracellular cytokine expression was assessed, cells were stimulated for 18 h with Aβ1-42 fibrils (1 μM final; rPeptide) in the presence of brefeldin A (2 μg/mL; Sigma-Aldrich) prior to staining. Data were acquired on an LSRFortessa flow cytometer at the Cincinnati Children’s Hospital Flow Cytometry Core and analyzed using FlowJo software (version 10.8.1). Fluorescence-minus-one controls were used when appropriate to establish gating boundaries.

### 2.7 Brain processing and histology

For histological analysis, brains were collected at the 9-week time point after the final treatment injection, following completion of behavioral testing.

#### Brain tissue processing and sectioning

Mice were transcardially perfused with 30 mL ice-cold PBS, and brains were harvested and fixed in 10% neutral buffered formalin for at least 24 h. After cryoprotection in 20% sucrose for 24 h and 30% sucrose for 48 h, brains were embedded in OCT, frozen on dry ice, and sectioned at 30 μm using a cryostat at the University of Cincinnati College of Medicine Microscopy Core. For brains collected from mice injected with FITC-PLGA-IMPs, samples were protected from direct light throughout fixation and sectioning.

#### Thioflavin-S staining

Free-floating brain sections were washed in PBS and dehydrated through graded ethanol solutions (100%, 90%, 70%, and 50%) for 3 min each. Sections were then incubated in 0.5% Thioflavin-S (Sigma) in 50% ethanol for 10 min, washed in PBS, and mounted with histology mounting medium.

#### Fluoro-Jade C staining

Neurodegeneration was assessed using a Fluoro-Jade C staining kit (Biosensis, TR-100-FJT). Free-floating sections were washed in PBS and incubated for 10 min in a staining solution containing Fluoro-Jade C and DAPI diluted in distilled water. Sections were then washed in PBS, incubated in 50 mM ammonium chloride for 15 min, washed again in PBS, and mounted with histology mounting medium.

#### Immunostaining

For immunofluorescence staining, free-floating sections were fixed in acetone at −20°C for 10 min and dehydrated in 70% ethanol for 5 min at −20°C. Sections were washed in PBS containing 0.05% Tween-20, blocked in 10% donkey serum for 30 min, and incubated with primary antibodies diluted in 3% donkey serum for 1 h at room temperature. The following primary antibodies were used: rabbit anti-Iba1 (FUJIFILM Wako, 1:1000) and rat anti-mouse CD68 (Bio-Rad, 1:200). After washing, sections were incubated with donkey anti-rabbit or donkey anti-rat secondary antibodies (Invitrogen, 1:500) for 30 min, washed again, incubated in 1% Triton X-100 for 10 min, and mounted with VECTASHIELD mounting medium containing DAPI (Vector Laboratories).

### 2.8 Image acquisition and quantification

Images of stained brain sections were acquired using an EVOS M5000 microscope at 10x or 20x magnification. Quantification was performed on 10x images using ImageJ software (version 1.54g). Thresholds were set independently for each staining condition and then applied consistently across all images within that staining set to distinguish positive signal from background. For CD68 and Iba1 staining, particle size thresholds were set at 10-2500 pixels^2^. For Thioflavin-S and Fluoro-Jade C staining, particle size thresholds were set at 50-2500 pixels^2^. Circularity was set at 0.0-1.0 for all analyses. Each data point represents one mouse and is expressed as the average number of positive signals across 3-4 brain sections.

### 2.9 Statistical analysis

Data are presented as mean ± SEM. Statistical significance was determined using one-way ANOVA followed by Tukey’s multiple-comparisons test, unpaired two-tailed Student’s *t* test, or two-sided Fisher’s exact test, as appropriate. Statistical analyses were performed using GraphPad Prism version 11.0. Differences were considered statistically significant at *p* < 0.05.

## 3. Results

### 3.1 Physicochemical characterization of PLGA-IMPs and FITC-PLGA-IMPs

To generate immunomodulatory poly(lactic-co-glycolic acid) immune-modifying particles (PLGA-IMPs), nanoparticles were synthesized in batches using a single-emulsion solvent evaporation protocol adapted from previously published methods [37, 48]. Because particle size, surface charge and polydispersity are key determinants of nanoparticle behavior, these properties were assessed by dynamic light scattering (DLS) and Zeta-potential analysis. Representative DLS profiles are shown in **Fig. 1A**. Across independent preparations, PLGA-IMPs displayed a Z-average diameter of 515.74 ± 42.92 nm, a Zeta-potential of -72.03 ± 1.60 mV, and a polydispersity index (PdI) of 0.216 ± 0.043 (**Fig. 1B**). Although a small secondary peak was occasionally detected in representative traces, it contributed less than 1% of the total intensity and was therefore considered negligible relative to the dominant particle population, supporting reproducible generation of particles with the desired size range and negative surface charge.

**Figure 1.**
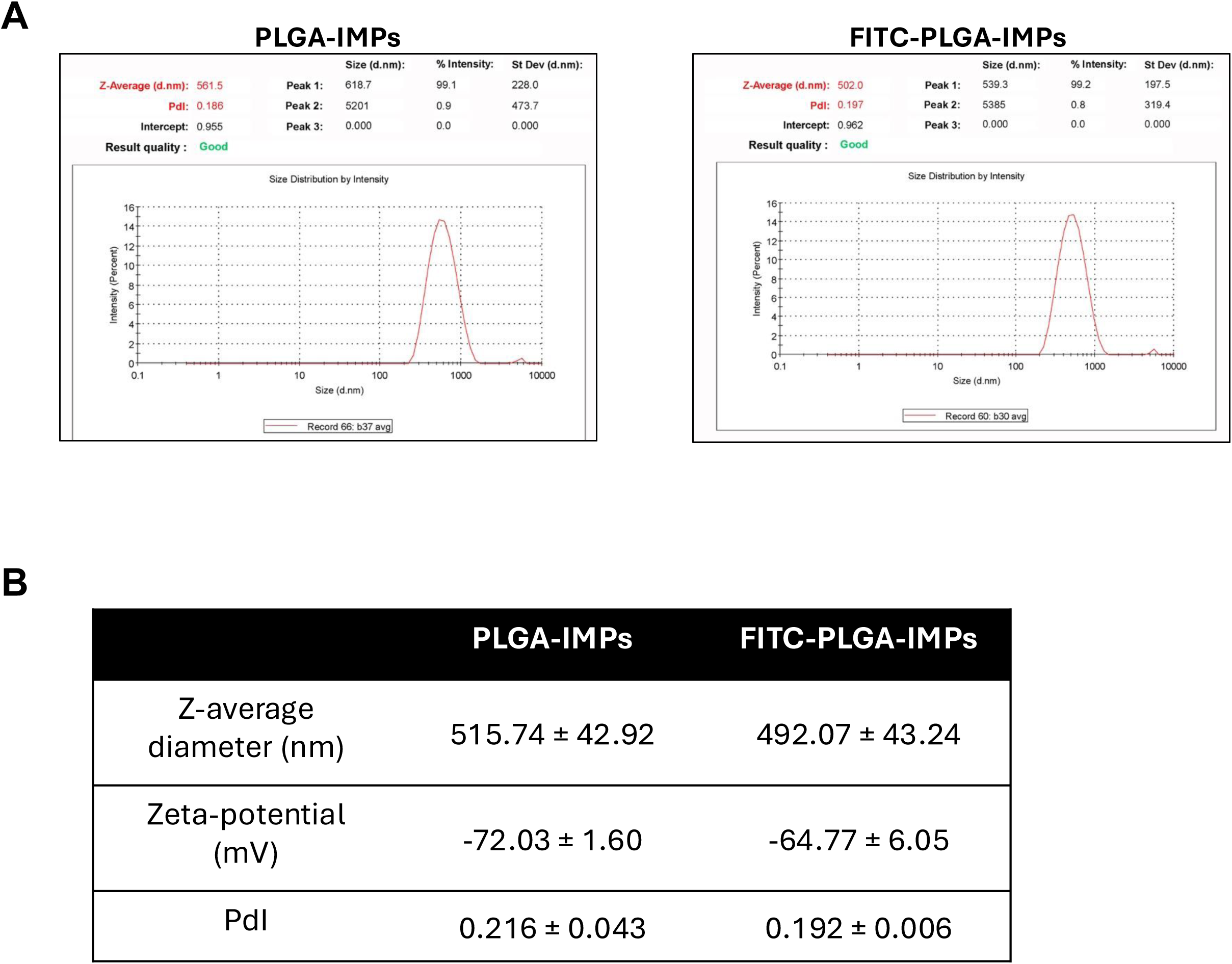
Physicochemical characterization of PLGA-IMPs and FITC-PLGA-IMPs. **(A)** Representative dynamic light scattering (DLS) profiles of unlabeled PLGA-IMPs and FITC-PLGA-IMPs showing particle size distribution by intensity. **(B)** Quantification of Z-average diameter, Zeta-potential, and polydispersity index (PdI) for PLGA-IMPs and FITC-PLGA-IMPs. Values are presented as mean ± SEM from independent particle preparations (PLGA-IMPs, *n* = 5; FITC-PLGA-IMPs, *n* = 3). FITC labeling did not substantially alter the physicochemical properties of the particles.

To track particle uptake, fluorescent PLGA-IMPs were generated using FITC-conjugated PLGA and formulated using the same protocol as unlabeled particles. FITC-PLGA-IMPs showed physicochemical properties comparable to those of unlabeled PLGA-IMPs, with a Z-average diameter of 492.07 ± 43.24 nm, a Zeta-potential of -64.77 ± 6.05 mV, and a PdI of 0.192 ± 0.006 (**Fig. 1A** and **1B**). These data indicate that FITC labeling did not substantially alter the size distribution, polydispersity, or surface charge of the particles.

*3.2 PLGA-IMPs are preferentially taken up by myeloid cells in the blood, spleen, and brain of WT and 5xFAD mice* To assess the *in vivo* cellular distribution of PLGA-IMPs, FITC-labeled PLGA-IMPs were administered to WT and 5xFAD mice, and immune cells were isolated from the blood, spleen, and brain 6 h later for flow cytometric analysis. WT and 5xFAD mice were used to compare particle uptake in healthy and disease-associated inflammatory environments.

In both WT and 5xFAD mice, FITC-PLGA-IMPs were taken up predominantly by CD11b^+^ myeloid cells rather than CD3^+^ T cells in the blood and spleen (**Fig. 2A**). Uptake by CD11b^+^ cells was similar between WT and 5xFAD mice in these peripheral compartments, whereas CD3^+^ T cells showed minimal particle association in both groups (**Fig. 2A**).

**Figure 2.**
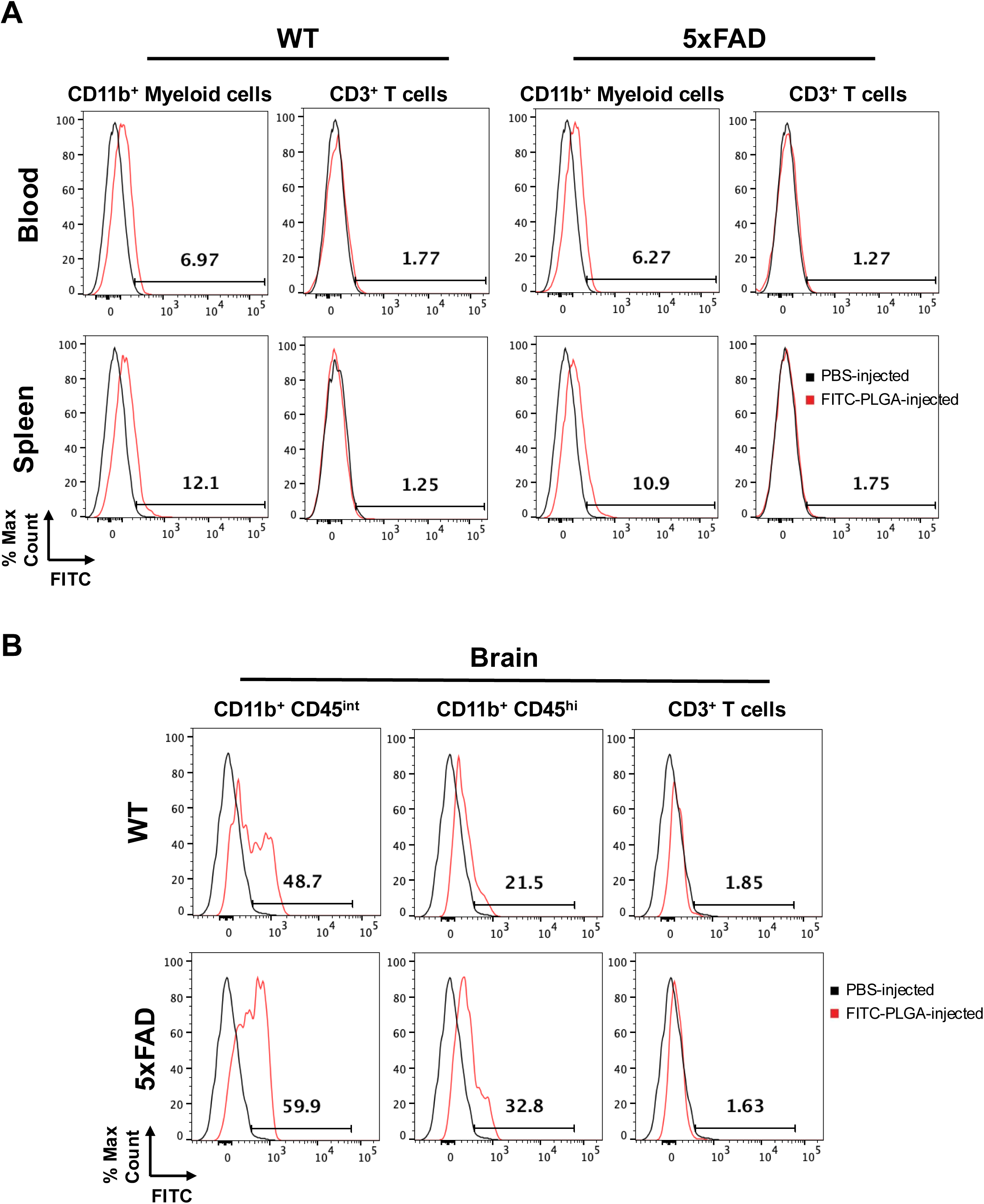
FITC-labeled PLGA-IMPs preferentially associate with myeloid cells in the blood, spleen, and brain of WT and 5xFAD mice. WT and 5xFAD mice (6 months old) received an intraperitoneal injection of FITC-labeled PLGA-IMPs (2.5 mg), and immune cells were isolated from the blood, spleen, and brain 6 h later for flow cytometric analysis. **(A)** Representative histograms showing FITC-PLGA-IMP association with CD11b^+^ myeloid cells and CD3^+^ T cells in the blood and spleen of WT and 5xFAD mice. **(B)** Representative histograms showing FITC-PLGA-IMP association with CD45^hi^CD11b^+^ and CD45^int^CD11b^+^ brain myeloid cell populations, as well as CD3^+^ T cells, in WT and 5xFAD mice. Black histograms represent tissue-matched background controls from PBS-injected mice, and red histograms represent cells isolated 6 h after FITC-PLGA-IMP injection. Numbers indicate the percentage of FITC-positive cells within each gated population. Data shown are representative of *n* = 2 independent experiments.

We next examined whether PLGA-IMPs could also be detected in brain immune populations. The gating strategy used to define the brain populations analyzed in these experiments is shown in **Fig. S1**. In the brain, both CD45^hi^CD11b^+^ and CD45^int^CD11b^+^ myeloid populations showed markedly greater FITC-PLGA-IMP uptake than CD3^+^ T cells (**Fig. 2B**). Notably, the proportion of FITC-positive CD45^int^CD11b^+^ and CD45^hi^CD11b^+^ cells was higher in the brains of 5xFAD mice than in WT mice (**Fig. 2B**). In addition, uptake by brain myeloid populations was greater than that observed in blood or splenic myeloid cells. Together, these findings indicate that PLGA-IMPs preferentially associate with myeloid cells *in vivo*, suggesting that uptake by brain myeloid populations is increased in the 5xFAD disease setting.

### 3.3 PLGA-IMP treatment ameliorates behavioral abnormalities in 5xFAD mice

To evaluate the therapeutic potential of PLGA-IMPs after disease onset, 6.5-month-old 5xFAD mice were treated with PLGA-IMPs or PBS three times per week for 4 weeks. Behavioral testing was then performed at 3 weeks and 9 weeks after the final injection to assess both early and sustained treatment effects. Elevated plus maze testing was used to assess exploratory and risk-associated behavior, whereas Barnes maze testing was used to evaluate spatial learning and memory. For the 9-week Barnes maze assessment, the escape hole was moved to the opposite quadrant to minimize bias from the initial training phase.

At both 3 weeks and 9 weeks after treatment, PBS-treated 5xFAD mice spent more time in the open arms of the elevated plus maze than WT controls, consistent with prior reports showing increased open-arm exploration in 5xFAD mice across multiple ages [10] (**Fig. 3A** and **3B**). In contrast, PLGA-IMP-treated 5xFAD mice spent significantly less time in the open arms than PBS-treated 5xFAD mice, with values shifted toward those observed in WT mice (**Fig. 3A** and **3B**). Total distance traveled did not differ significantly between PLGA-IMP- and PBS-treated 5xFAD mice at either time point, indicating that the change in open-arm behavior was not attributable to altered locomotor activity (**Fig. 3A** and **3B**).

**Figure 3.**
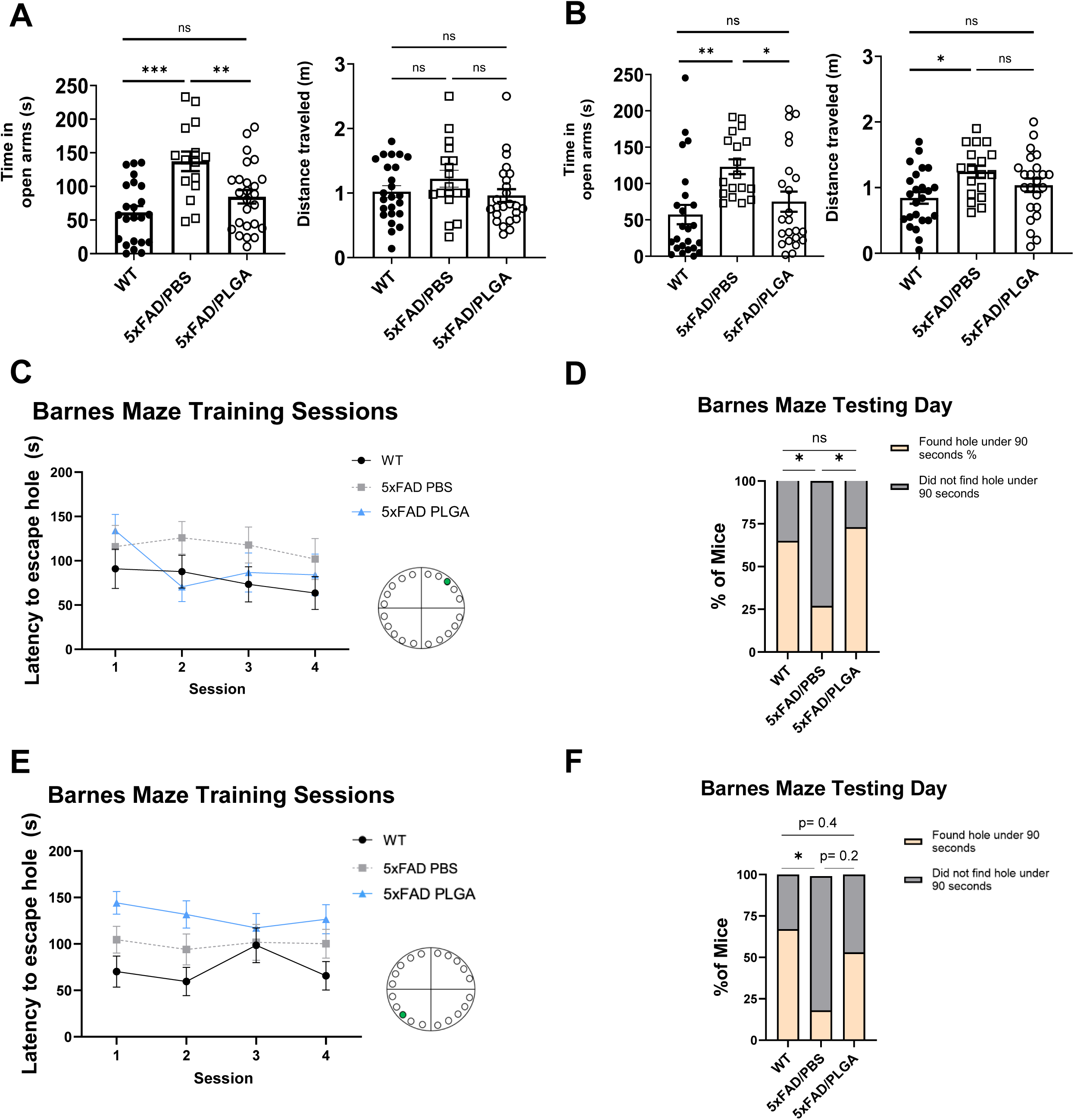
PLGA-IMP treatment attenuates behavioral abnormalities in 5xFAD mice. Behavioral analysis of WT, 5xFAD/PBS, and 5xFAD/PLGA mice was performed using the elevated plus maze (EPM) and Barnes maze at 3 weeks (A, C, D) and 9 weeks (B, E, F) after the final treatment injection. (**A**) Time spent in the open arms and total distance traveled during the 5-min EPM test at 3 weeks after treatment in WT (n = 24), 5xFAD/PBS (n = 18), and 5xFAD/PLGA (n = 24) mice. (**B**) Time spent in the open arms and total distance traveled during the 5-min EPM test at 9 weeks after treatment in WT (n = 24), 5xFAD/PBS (n = 18), and 5xFAD/PLGA (n = 23) mice. (**C**) Barnes maze training performance at 3 weeks after treatment, shown as latency to locate the escape hole across 4 training sessions in WT (n = 13), 5xFAD/PBS (n = 11), and 5xFAD/PLGA (n = 10) mice. The location of the escape hole is indicated in green. (**D**) Percentage of mice that located the escape hole within 90 s on the Barnes maze test day at 3 weeks after treatment in WT (n = 13), 5xFAD/PBS (n = 11), and 5xFAD/PLGA (n = 10) mice. (**E**) Barnes maze training performance at 9 weeks after treatment, shown as latency to locate the escape hole across 4 training sessions in WT (n = 17), 5xFAD/PBS (n = 17), and 5xFAD/PLGA (n = 18) mice. The location of the escape hole was moved to the opposite quadrant relative to the 3-week assessment and is indicated in green. (**F**) Percentage of mice that located the escape hole within 90 s on the Barnes maze test day at 9 weeks after treatment in WT (n = 13), 5xFAD/PBS (n = 11), and 5xFAD/PLGA (n = 15) mice. Data are presented as mean ± SEM in A-C and E. In A and B, each symbol represents one mouse. \**p* < 0.05, ** *p* < 0.01, *** *p* < 0.001; ns, not significant. Panels A and B were analyzed by one-way ANOVA with multiple-comparisons testing. Panels D and F were analyzed by Fisher’s exact test. Statistical comparisons for panels C and E were performed using session 4 data.

In the Barnes maze, escape latency during training sessions did not reveal clear group differences at either 3 weeks or 9 weeks after treatment (**Fig. 3C** and **3E**), suggesting that PLGA-IMP treatment did not markedly alter acquisition during the training phase. However, on the testing day, a greater proportion of PLGA-IMP-treated 5xFAD mice located the escape hole within 90 s compared with PBS-treated 5xFAD mice at 3 weeks after treatment (**Fig. 3D**). A similar trend was observed at 9 weeks, although this did not reach statistical significance (**Fig. 3F**). Together, these data indicate that PLGA-IMP treatment ameliorates behavioral abnormalities in 5xFAD mice, with a robust effect in the elevated plus maze and a more modest effect in Barnes maze performance.

### 3.4 PLGA-IMP treatment induces persistent changes in brain microglial populations 9 weeks after treatment

To determine whether PLGA-IMP treatment induced durable changes in brain immune populations, brains were collected from WT, PBS-treated 5xFAD, and PLGA-IMP-treated 5xFAD mice immediately after behavioral testing at the 9-week time point and analyzed by flow cytometry. A representative gating strategy, including the CD11c fluorescence-minus-one control used to define the P2RY12^+^CD11c^+^ population, is shown in **Fig. S2**. Compared with WT mice, PBS-treated 5xFAD mice exhibited increased frequencies of total CD45^+^ leukocytes, CD45^+^CD11b^-^ lymphoid cells, CD11b^+^P2RY12^-^ myeloid cells, and P2RY12^+^CD11c^+^ microglia, together with a reduction in the proportion of CD11b+P2RY12+ microglia (**Fig. 4A**). PLGA-IMP treatment significantly reduced the frequencies of total CD45^+^ cells, CD45^+^CD11b^-^ lymphoid cells, CD11b^+^P2RY12^-^ myeloid cells, and P2RY12^+^CD11c^+^ microglia in 5xFAD brains, but did not restore the overall proportion of CD11b^+^P2RY12^+^ microglia (**Fig. 4A**).

**Figure 4.**
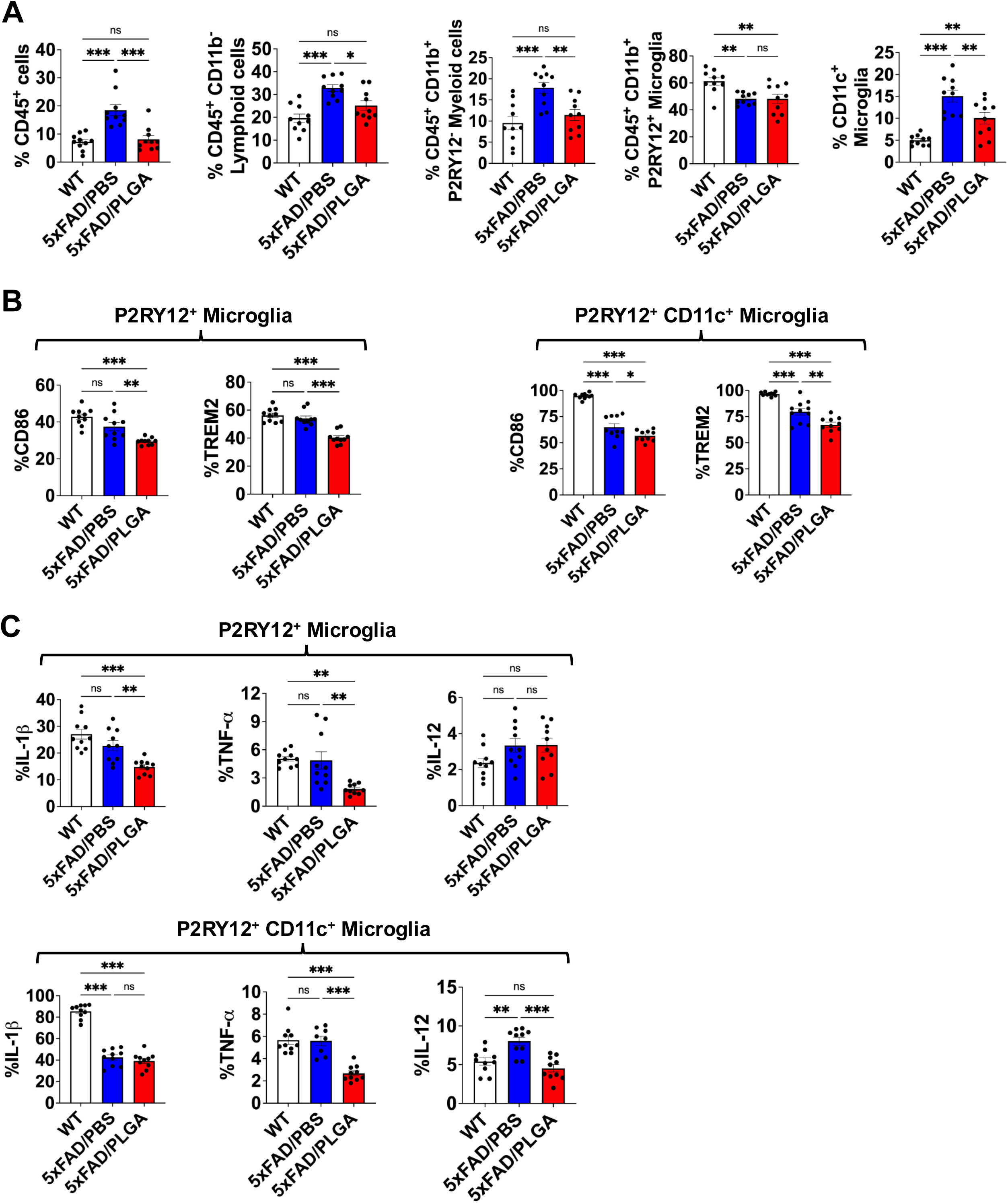
PLGA-IMP treatment induces persistent changes in brain immune composition and microglial phenotype in 5xFAD mice 9 weeks after treatment. (**A**) Flow cytometric analysis of brain immune composition in WT, 5xFAD/PBS, and 5xFAD/PLGA mice at 9 weeks after the final treatment injection, showing frequencies of total CD45^+^ leukocytes, CD45^+^CD11b^-^ lymphoid cells, CD45^+^CD11b^+^P2RY12^-^ myeloid cells, CD45^+^CD11b^+^P2RY12^+^ microglia, and P2RY12^+^CD11c^+^ microglia. (**B**) Quantification of CD86^+^ and TREM2^+^ cells within P2RY12^+^ microglia and P2RY12^+^CD11c^+^ microglia. (**C**) Frequencies of IL-1β^+^, TNF-α^+^, and IL-12p40^+^ cells within P2RY12^+^ microglia and P2RY12^+^CD11c^+^ microglia. Data are presented as mean ± SEM, with each symbol representing one mouse (*n* = 10 mice per group). Statistical significance was determined by one-way ANOVA followed by Tukey’s multiple-comparisons test. \**p* < 0.05, ** *p* < 0.01, *** *p* < 0.001; ns, not significant.

We next examined activation- and disease-associated markers within microglial populations. Within the P2RY12^+^ microglial compartment, PLGA-IMP treatment reduced the frequencies of CD86^+^ and TREM2^+^ cells compared with PBS-treated 5xFAD mice (**Fig. 4B**). PLGA-IMP treatment also reduced the frequencies of IL-1β^+^ and TNF-α^+^ P2RY12^+^ microglia, whereas IL-12p40 expression was not significantly altered (**Fig. 4C**). A similar but more pronounced pattern was observed within the P2RY12^+^CD11c^+^ microglial subset. In this population, PLGA-IMP treatment reduced CD86 and TREM2 expression and was associated with lower TNF-α and IL-12p40 expression compared with PBS-treated 5xFAD mice, whereas IL-1β remained elevated and was not significantly changed by treatment (**Fig. 4B** and **4C**). IL-10 and CD80 analysis revealed no differences (data not shown).

By contrast, analysis of CD11b^+^P2RY12^-^ myeloid cells revealed limited evidence of a sustained treatment effect at this time point. Although several markers differed between WT and 5xFAD mice, CD86, TREM2, IL-1β, TNF-α, and IL-12p40 expression did not differ significantly between PBS- and PLGA-IMP-treated 5xFAD groups (**Fig. S3**). Together, these findings indicate that PLGA-IMP treatment induces persistent changes in brain immune composition and predominantly modulates resident microglial populations, particularly CD11c^+^ microglia, with comparatively limited sustained effects in CD11b^+^P2RY12^-^ myeloid cells at 9 weeks after treatment.

### 3.5 PLGA-IMP treatment reduces key pathological hallmarks of AD in the brains of 5xFAD mice

To determine whether the behavioral effects of PLGA-IMP treatment were accompanied by changes in brain pathology, we examined amyloid plaque burden, CD68 immunoreactivity, and neurodegeneration in the cortex and hippocampus of WT and 5xFAD mice 9 weeks after the final treatment. These regions were selected because both are prominently affected in 5xFAD mice and are relevant to the behavioral abnormalities assessed in this study.

Thioflavin-S staining revealed that PLGA-IMP-treated 5xFAD mice exhibited reduced amyloid plaque burden in the cortex compared with PBS-treated 5xFAD mice (**Fig. 5A** and **5B**). In contrast, plaque burden in the hippocampus was not significantly different between treatment groups (**Fig. 5B**).

**Figure 5.**
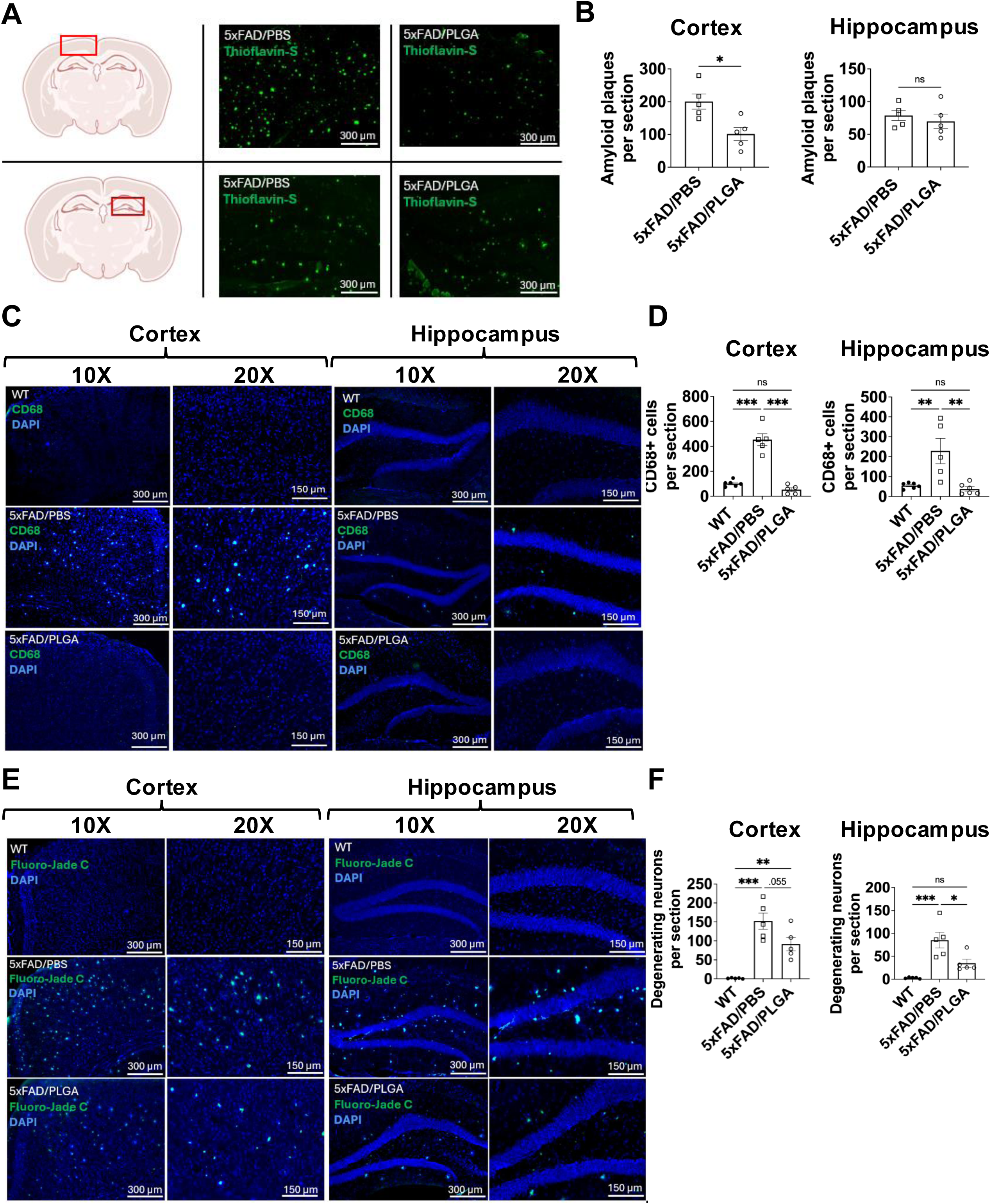
PLGA-IMP treatment reduces amyloid plaque burden, CD68 immunoreactivity, and neurodegeneration in the brains of 5xFAD mice. (**A**) Representative Thioflavin-S staining of amyloid plaques in the cortex and hippocampus of 5xFAD mice 9 weeks after treatment with PBS or PLGA-IMPs. Red boxes on the schematic indicate the analyzed regions. Scale bars, 300 μm. (**B**) Quantification of Thioflavin-S+ amyloid plaques in the cortex (left panel) and hippocampus (right panel) of 5xFAD/PBS (n = 5) and 5xFAD/PLGA (n = 5) mice. (**C**) Representative CD68 immunofluorescence in the cortex and hippocampus of WT, 5xFAD/PBS, and 5xFAD/PLGA mice 9 weeks after treatment. Nuclei were counterstained with DAPI. Scale bars, 300 μm and 150 μm as indicated. (**D**) Quantification of CD68+ cells in the cortex (left panel) and hippocampus (right panel) of WT (n = 5), 5xFAD/PBS (n = 5), and 5xFAD/PLGA (n = 5) mice. (**E**) Representative Fluoro-Jade C staining of degenerating neurons in the cortex and hippocampus of WT, 5xFAD/PBS, and 5xFAD/PLGA mice 9 weeks after treatment. Nuclei were counterstained with DAPI. Scale bars, 300 μm and 150 μm as indicated. (**F**) Quantification of Fluoro-Jade C+ degenerating neurons in the cortex (left panel) and hippocampus (right panel) of WT (n = 5), 5xFAD/PBS (n = 5), and 5xFAD/PLGA (n = 5) mice. Data are presented as mean ± SEM, with each symbol representing one mouse. Each brain region was analyzed separately. \**p* < 0.05, ** *p* < 0.01, *** *p* < 0.001; ns, not significant. Panel B was analyzed using an unpaired two-tailed Student’s t-test, whereas panels D and F were analyzed using one-way ANOVA followed by Tukey’s multiple-comparisons test.

We next assessed CD68 immunoreactivity as a marker associated with activated myeloid cells. As expected, PBS-treated 5xFAD mice showed a marked increase in CD68+ cells compared with WT mice, particularly in the cortex (**Fig. 5C** and **5D**). PLGA-IMP treatment significantly reduced the number of CD68+ cells in the cortex, with a similar trend observed in the hippocampus (**Fig. 5C** and **5D**).

Finally, Fluoro-Jade C staining was used to assess neurodegeneration. PBS-treated 5xFAD mice showed increased numbers of degenerating neurons in both cortex and hippocampus relative to WT mice, whereas PLGA-IMP-treated 5xFAD mice exhibited fewer Fluoro-Jade C+ cells in both regions (**Fig. 5E** and **5F**). This reduction was significant in the hippocampus and showed a similar trend in the cortex (**Fig. 5F**). Additional representative histological fields corresponding to these analyses are shown in **Fig. S4**.

Together, these data indicate that PLGA-IMP treatment is associated with reduced cortical plaque burden, decreased CD68 immunoreactivity, and reduced neurodegeneration in 5xFAD mice, supporting a sustained beneficial effect on multiple pathological features of disease.

## 4. Discussion

In this study, we show that systemically administered PLGA-IMPs produce durable beneficial effects in the 5xFAD model of amyloid-driven neurodegeneration. Treatment of 6.5-month-old 5xFAD mice with intraperitoneal PLGA-IMPs was associated with altered *in vivo* particle uptake by myeloid populations, improvement in behavioral abnormalities, persistent changes in brain immune composition, and reduction of multiple pathological hallmarks of disease, including cortical amyloid plaque burden, CD68 immunoreactivity, and neurodegeneration. Together, these findings support the therapeutic potential of PLGA-IMPs in AD and suggest that their effects are mediated, at least in part, through long-lasting modulation of brain myeloid responses.

An important feature of this study is the timing and route of treatment. Prior work with native PLGA in AD-related models showed beneficial effects on amyloid pathology and behavior, but that study used intracerebroventricular delivery in young, 3-month-old 5xFAD mice [1]. In contrast, our study tested a less invasive systemic route and initiated treatment after behavioral abnormalities were already established. Thus, the present work extends prior observations by showing that PLGA-based particles can remain effective when administered intraperitoneally at a later, more therapeutically relevant stage of disease.

The behavioral data suggest that PLGA-IMPs exert both early and persistent functional benefits. The most robust changes were observed in the elevated plus maze, where PLGA-IMP-treated 5xFAD mice consistently spent less time in the open arms than PBS-treated 5xFAD mice at both 3 and 9 weeks after treatment, without changes in total distance traveled. This indicates that the reduction in open-arm time was not simply due to reduced overall locomotor activity. Increased open-arm exploration in 5xFAD mice has been reported previously [23] and has been interpreted as altered exploratory or risk-associated behavior rather than a straightforward reduction in anxiety [8]. One possible explanation is that this phenotype reflects circuit dysfunction secondary to degeneration or impaired function of inhibitory neuronal populations [8, 14], which has been linked to cortical and hippocampal hyperexcitability in this model [31]. Within that framework, the improvement observed after PLGA-IMP treatment may reflect partial preservation of neural circuit integrity resulting from reduced neuroinflammation and neurodegeneration. The Barnes maze findings were more modest, but still informative. PLGA-IMP treatment did not clearly alter acquisition during training, yet more treated mice located the escape hole within 90 s on the test day, particularly at the earlier post-treatment time point. The decision to move the escape hole to the opposite quadrant for the 9-week assessment is important, because it reduces simple carryover from the initial training phase and places greater emphasis on updated spatial learning and memory retrieval. The more modest effect observed in the Barnes maze compared with the elevated plus maze may therefore reflect the fact that these tasks probe distinct neural processes and differ in their sensitivity to inflammatory and structural changes across disease progression.

One of the most interesting findings of the study is that the persistent immune effect of PLGA-IMPs at 9 weeks after treatment was strongest in P2RY12^+^ microglial populations, particularly the CD11c^+^ subset, rather than in CD11b^+^P2RY12^-^ myeloid cells. PBS-treated 5xFAD mice showed increased total leukocyte infiltration, expansion of CD11c^+^ microglia, and altered expression of activation- and disease-associated markers within microglial populations. PLGA-IMP treatment reduced several of these abnormalities and was associated with lower CD86, TREM2, IL-1β, and TNF-α expression within microglial subsets, whereas the CD11b^+^P2RY12^-^ population showed little sustained treatment responsiveness. These data suggest that although PLGA-IMPs are preferentially associated with myeloid cells *in vivo*, their long-term impact in the AD brain may be expressed primarily through persistent reshaping of microglial states. This interpretation is also consistent with the uptake experiment shown in Fig. 2, in which FITC-PLGA-IMPs were most strongly associated with CD45^int^CD11b^+^ brain myeloid cells, a population consistent with resident microglia.

This interpretation is especially relevant in light of the growing literature on CD11c^+^ microglia in AD. CD11c was included in our analysis because it has been widely associated with disease-associated microglial (DAM) [25, 27] and plaque-associated microglial (PAM) states [43], and prior work indicates that CD11c^+^ microglia substantially overlap with transcriptionally defined disease microglial programs [27, 47]. At the same time, CD11c is best viewed as a useful but imperfect marker, as CD11c^+^ cells are heterogeneous and their functional significance depends on molecular context and disease stage [13, 43]. Indeed, prior studies have supported both protective [27] and pathogenic [33, 43] roles for CD11c^+^ microglia, and recent work indicates that this compartment itself contains functionally distinct subsets rather than representing a uniformly beneficial or uniformly harmful population [43, 60]. In this study, the reduction in P2RY12^+^CD11c^+^ microglia after PLGA-IMP treatment is therefore most appropriately interpreted as modulation of a disease-associated microglial state rather than elimination of a uniformly harmful population.

A similar caution applies to TREM2, which is closely linked to the biology of these disease-associated microglial states. TREM2 is a central regulator of microglial responses to amyloid pathology and is required for full induction of the DAM program, with many studies supporting roles in plaque association, phagocytosis, metabolism, and neuroprotection [7, 15, 27, 39, 49, 52, 57]. However, TREM2 is also embedded within a broader and stage-dependent disease-associated program, and increased TREM2 expression should not be assumed to reflect a uniformly beneficial state in all contexts. In the present study, the decrease in TREM2 expression observed after PLGA-IMP treatment is therefore best interpreted not as direct inhibition of a protective pathway, but as part of a broader remodeling of a disease-associated microglial phenotype at a late post-treatment time point. It is also possible that lower TREM2 expression reflects reduced ongoing engagement of microglia with amyloid pathology, consistent with the reduced plaque burden observed in treated mice. Together, the CD11c and TREM2 findings support the view that PLGA-IMPs do not merely suppress microglial activation globally but instead reshape a specific disease-associated microglial compartment that is likely functionally heterogeneous. Thus, our findings should not be interpreted as evidence that therapeutic suppression of TREM2 is beneficial in AD, but rather that successful modulation of disease-associated microglial states may secondarily reduce TREM2 expression once pathological burden is diminished.

One unresolved question is how these long-lasting changes in resident microglial states are established. One possibility is that PLGA-IMPs alter the inflammatory milieu early after administration, thereby contributing to reduced accumulation of peripheral immune cells in the CNS during a critical phase of ongoing pathology. Even if this effect is transient, it could secondarily influence resident microglia and promote a more durable shift in the brain immune environment. This model may help explain the apparent longevity of treatment. Such an interpretation would be consistent with prior work showing that negatively charged immune-modifying particles can modulate inflammatory myeloid-cell behavior in CNS injury and inflammatory disease models [11, 24, 40, 45]. In the present study, the fact that the strongest late effects were observed in P2RY12^+^ microglia rather than in CD11b^+^P2RY12^-^ myeloid cells supports the idea that PLGA-IMPs may initiate a cascade of immune remodeling whose most persistent effects are retained within resident brain myeloid populations.

The physicochemical properties of the particles may be relevant to this biology. The PLGA-IMP batches used here displayed diameters in the roughly 400-700 nm range and a negative surface charge, and FITC-labeled particles retained similar properties. Particles in this size range are expected to require active phagocytic uptake rather than passive membrane diffusion [16, 35], which likely contributes to the preferential association with myeloid cells seen in the blood, spleen, and brain. In this sense, the size of the particles is not simply a manufacturing feature, but part of the therapeutic design, favoring uptake by professional phagocytes and thereby biasing the response toward innate immune populations.

Several non-mutually exclusive mechanisms could account for the downstream biological effects of PLGA-IMPs. Because these particles are large, negatively charged, and taken up by phagocytic cells, they may mimic some aspects of efferocytic cargo or cell debris and thereby alter inflammatory programming after uptake [9, 36, 44]. It is also possible that PLGA directly interacts with extracellular Aβ species, as suggested by prior work using native PLGA, which may contribute to reduced plaque burden [1]. However, our data do not directly test either of these possibilities. Similarly, although the degradation products of PLGA can enter endogenous metabolic pathways and may influence cellular homeostasis, the present study does not establish a mitochondrial or metabolic mechanism. For this reason, we think the most appropriate interpretation is that PLGA-IMPs act through a combination of physicochemical targeting to phagocytic myeloid cells and subsequent immunomodulatory effects, rather than through one clearly defined pathway alone.

The pathology data further support this model. PLGA-IMP treatment reduced cortical amyloid plaque burden, decreased CD68 immunoreactivity, and reduced neurodegeneration, particularly in the hippocampus. These findings are important because they link the immune changes identified by flow cytometry to tissue-level outcomes that are relevant to cognitive decline. The regional pattern was not identical across all readouts, with plaque reduction being strongest in cortex and neuroprotection more evident in hippocampus. This likely reflects the fact that amyloid deposition, myeloid activation, and neuronal vulnerability do not progress uniformly across brain regions in 5xFAD mice. Nevertheless, the overall pattern supports the conclusion that PLGA-IMP treatment exerts a sustained protective effect on multiple disease-relevant features.

An additional strength of PLGA-IMPs is their translational attractiveness. Unlike more invasive delivery approaches, the present study used intraperitoneal administration. Moreover, PLGA is a biodegradable and well-established material with a strong safety record, and PLGA-based formulations have already been evaluated clinically in other disease settings. This is especially relevant in AD, where some current disease-modifying approaches remain limited by route, tolerability, or adverse effects. While our data do not establish clinical efficacy, they support the concept that a myeloid-targeted, biodegradable particle platform may offer a comparatively safe and scalable alternative or adjunctive approach for modifying neuroinflammatory aspects of disease.

This study also has limitations. The 5xFAD model captures robust amyloid pathology but does not model the full complexity of human AD, particularly tau pathology [38, 56]. Our mechanistic analyses were centered on myeloid populations and do not identify a single molecular pathway responsible for the observed benefit. In addition, some of the behavioral and histological effects were stronger in certain assays or regions than others, indicating that PLGA-IMP treatment does not uniformly normalize all disease features. Finally, although no sex-dependent differences were detected in the datasets analyzed here, the study was not specifically powered to define subtle sex-specific responses across all endpoints. These limitations notwithstanding, the convergent effects on behavior, microglial state, plaque burden, and neurodegeneration support the robustness of the overall therapeutic signal.

## 5. Conclusion

In summary, our study shows that systemically administered PLGA-IMPs produce durable beneficial effects in the 5xFAD model of Alzheimer’s disease. Treatment initiated after the onset of behavioral abnormalities improved exploratory/risk-associated behavior, reduced cortical amyloid plaque burden, decreased CD68 immunoreactivity, and attenuated neurodegeneration. These effects were accompanied by persistent changes in brain immune composition, with the most prominent long-term impact observed in resident microglial populations, particularly the CD11c^+^ subset, with comparatively limited sustained effects in CD11b^+^P2RY12^-^ myeloid cells. Together, these findings identify PLGA-IMPs as a promising biodegradable immunomodulatory platform capable of reshaping disease-associated myeloid responses in the Alzheimer’s disease brain. This work extends prior studies of native PLGA nanoparticles by demonstrating efficacy after systemic administration at a later, behaviorally symptomatic stage of disease, and supports further investigation of myeloid-targeted nanoparticle approaches as therapeutic strategies for Alzheimer’s disease.

## 6. Abbreviations

AD: Alzheimer’s disease
Aβ: amyloid beta
Aβ42: amyloid beta 42
APP: amyloid precursor protein
ARIA: amyloid-related imaging abnormalities
BACE1: beta-secretase 1
BSA: bovine serum albumin
CD: cluster of differentiation
CNS: central nervous system
DAM: disease-associated microglia
DAPI: 4′,6-diamidino-2-phenylindole
DI: deionized
DLS: dynamic light scattering
DMEM: Dulbecco’s Modified Eagle Medium
DNase I: deoxyribonuclease I
FBS: fetal bovine serum
FDA: Food and Drug Administration
FITC: fluorescein isothiocyanate
FMO: fluorescence-minus-one
GWAS: genome-wide association study
IACUC: Institutional Animal Care and Use Committee
Iba1: ionized calcium-binding adaptor molecule 1
IL-: interleukin
iNOS: inducible nitric oxide synthase i.p. intraperitoneal
iPSC: induced pluripotent stem cell
LAMS: Laboratory Animal Medical Services
NLRP3: NLR family pyrin domain containing 3
OCT: optimal cutting temperature
P2RY12: purinergic receptor P2Y12
PAM: plaque-associated microglia
PBS: phosphate-buffered saline
PdI: polydispersity index
PEMA: poly(ethylene-alt-maleic anhydride)
PLGA: poly(lactic-co-glycolic acid)
PLGA-IMPs: poly(lactic-co-glycolic acid) immune-modifying particles
PSEN1: presenilin 1
RPMI-1640: Roswell Park Memorial Institute 1640 medium
ROS: reactive oxygen species
SEM: standard error of the mean
TCA: tricarboxylic acid
TNF-α: tumor necrosis factor alpha
TREM2: triggering receptor expressed on myeloid cells 2
WT: wild type
5xFAD: transgenic mouse model carrying five familial Alzheimer’s disease mutations

## 7. Declarations

### Ethics approval and consent to participate

All animal procedures were performed in accordance with institutional guidelines and were approved by the Institutional Animal Care and Use Committee (IACUC) of the University of Cincinnati (protocol no. 23-08-22-01)

### Consent for publication

Not applicable

### Availability of data and materials

All data generated or analyzed during this study are included in this published article and its supplementary information files. Additional raw datasets are available from the corresponding author on reasonable request.

### Author Contributions

I.I. and B.S. conceived the study and designed the experiments. B.S. performed the majority of the experiments. K.S.P. and M.K. formulated the PLGA-IMPs and performed dynamic light scattering analysis. B.S., K.S.P., and M.K. contributed to data interpretation and critically revised the manuscript. B.S. and I.I. drafted the manuscript and revised it for intellectual content. All authors read and approved the final manuscript.

## Acknowledgements

We thank the University of Cincinnati Laboratory Animal Medical Services (LAMS) for animal care support, the Luo laboratory at the University of Cincinnati for access to behavioral testing equipment, and the Cincinnati Children’s Hospital Flow Cytometry Core for technical assistance.

