## Supplemental Information for "Poly(lactic-co-glycolic acid) immunomodulatory nanoparticles attenuate neuroinflammation and Alzheimer’s disease-related pathology in 5xFAD mice"

#### SUPPLEMENTAL FIGURE LEGENDS

**Figure S1. Representative gating strategy for analysis of brain immune populations in FITC-PLGA-IMP uptake experiments.** Representative plots illustrating the gating strategy used for the analysis of brain immune populations shown in Figure 2B. Sequential gates were applied to identify total cells, singlets, live cells, $CD45^{int}CD11b^{+}$ myeloid cells, $CD45^{hi}CD11b^{+}$ myeloid cells and $CD45^{+}CD11b^{-}CD3^{+}$ T cells.

**Figure S2. Representative gating strategy for flow cytometric analysis of brain myeloid populations in treatment-response experiments. (A)** Representative plots illustrating the sequential gating strategy used for the analyses shown in Figure 4. Gates were applied to identify total cells, singlets, live cells, $CD45^{+}$ leukocytes, $CD11b^{+}$ myeloid cells, and $P2RY12^{+}$ microglia. (**B**) Representative CD11c fluorescence-minus-one (FMO) control and CD11c staining within the gated $P2RY12^{+}$ microglial population, used to define the $P2RY12^{+}CD11c^{+}$ microglial subset analyzed in Figure 4

**Figure S3. PLGA-IMP treatment has limited long-term effects on $CD11b^{+}P2RY12^{-}$ myeloid cells in the brains of 5xFAD mice. (A)** Quantification of CD86 and TREM2 expression within $CD45^{+}CD11b^{+}P2RY12^{-}$ myeloid cells from WT, 5xFAD/PBS, and 5xFAD/PLGA mice at 9 weeks after the final treatment injection. (**B**) Quantification of IL-1β, TNF-α, and IL-12p40 expression within $CD45^{+}CD11b^{+}P2RY12^{-}$ myeloid cells from WT, 5xFAD/PBS, and 5xFAD/PLGA mice at 9 weeks after the final treatment injection. Data are presented as mean ± SEM, with each symbol representing one mouse (n = 10 mice per group). Statistical significance was determined by one-way ANOVA followed by Tukey's multiple-comparisons test. *$p < 0.05$, ** $p < 0.01$, *** $p < 0.001$; ns, not significant.

**Figure S4. Additional representative histological fields from cortex and hippocampus of PBS- and PLGA-IMP-treated 5xFAD mice.** Additional representative images corresponding to the histological analyses shown in Figure 5 are presented for Thioflavin-S staining (top row), CD68 immunostaining (middle row), and Fluoro-Jade C staining (bottom row) in the cortex (left) and hippocampus (right) of 5xFAD mice treated with PBS or PLGA-IMPs. Each image represents a different mouse and is included to provide qualitative support for the quantified pathological analyses shown in Figure 5.

### Figure S1

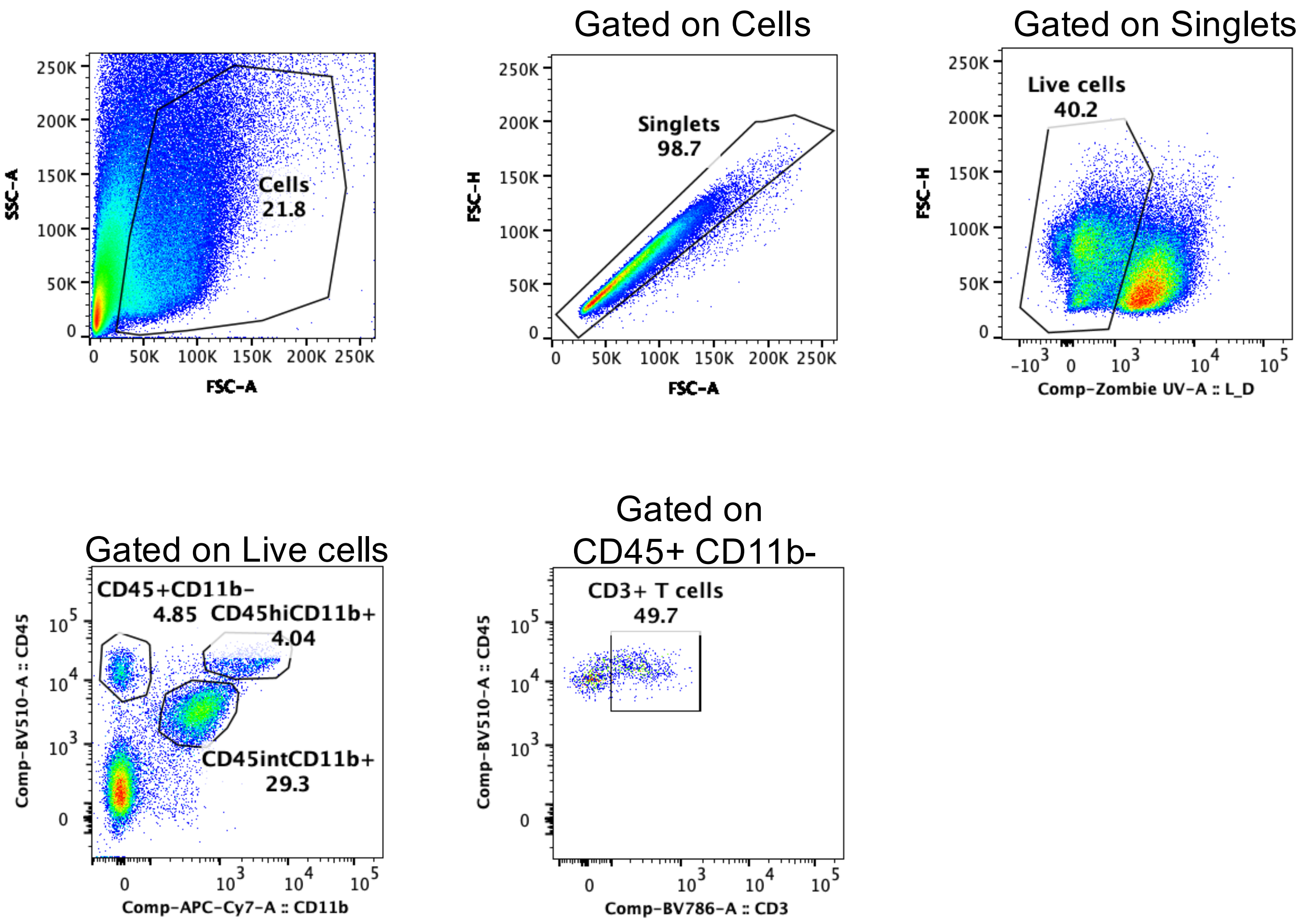

Gated on Cells
Gated on Singlets
Cells
21.8
SSC-A
FSC-A
Singlets
98.7
FSC-H
FSC-A
Live cells
40.2
FSC-H
Comp-Zombie UV-A :: L_D
Gated on Live cells
CD45+CD11b-
4.85
CD45hiCD11b+
4.04
CD45intCD11b+
29.3
Comp-BV510-A :: CD45
Comp-APC-Cy7-A :: CD11b
Gated on
CD45+ CD11b-
CD3+ T cells
49.7
Comp-BV510-A :: CD45
Comp-BV786-A :: CD3

### Figure S2

**A**

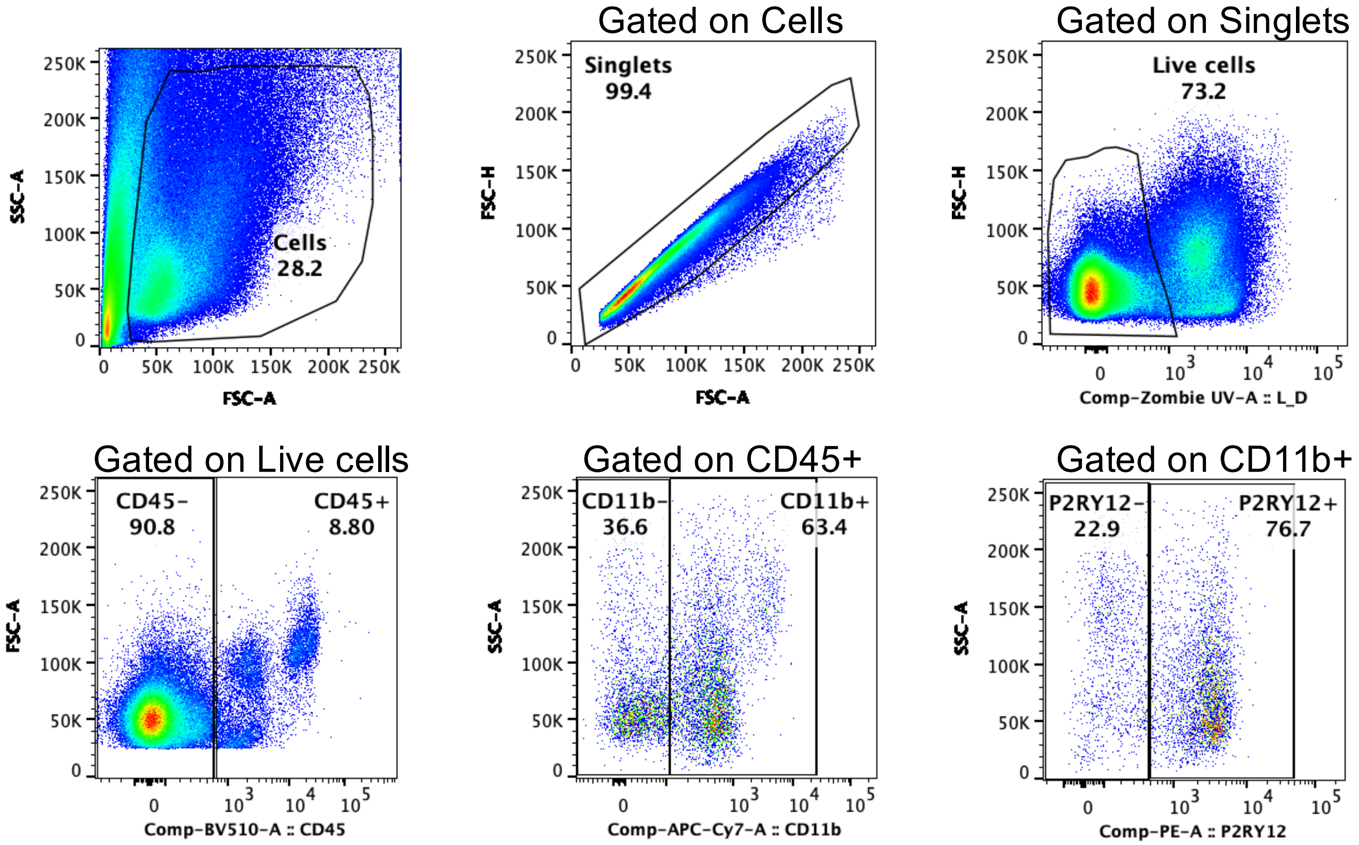

Cells
28.2
SSC-A
FSC-A
Gated on Cells
Singlets
99.4
FSC-H
FSC-A
Gated on Singlets
Live cells
73.2
FSC-H
Comp-Zombie UV-A :: L_D
Gated on Live cells
CD45-
90.8
CD45+
8.80
FSC-A
Comp-BV510-A :: CD45
Gated on CD45+
CD11b-
36.6
CD11b+
63.4
SSC-A
Comp-APC-Cy7-A :: CD11b
Gated on CD11b+
P2RY12-
22.9
P2RY12+
76.7
SSC-A
Comp-PE-A :: P2RY12


**B**

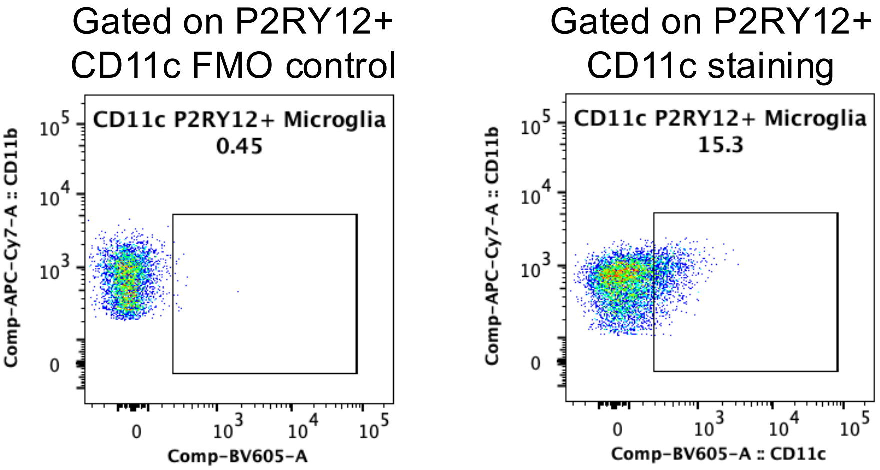

Gated on P2RY12+
CD11c FMO control
CD11c P2RY12+ Microglia
0.45
Comp-APC-Cy7-A :: CD11b
Comp-BV605-A
Gated on P2RY12+
CD11c staining
CD11c P2RY12+ Microglia
15.3
Comp-APC-Cy7-A :: CD11b
Comp-BV605-A :: CD11c

### Figure S3

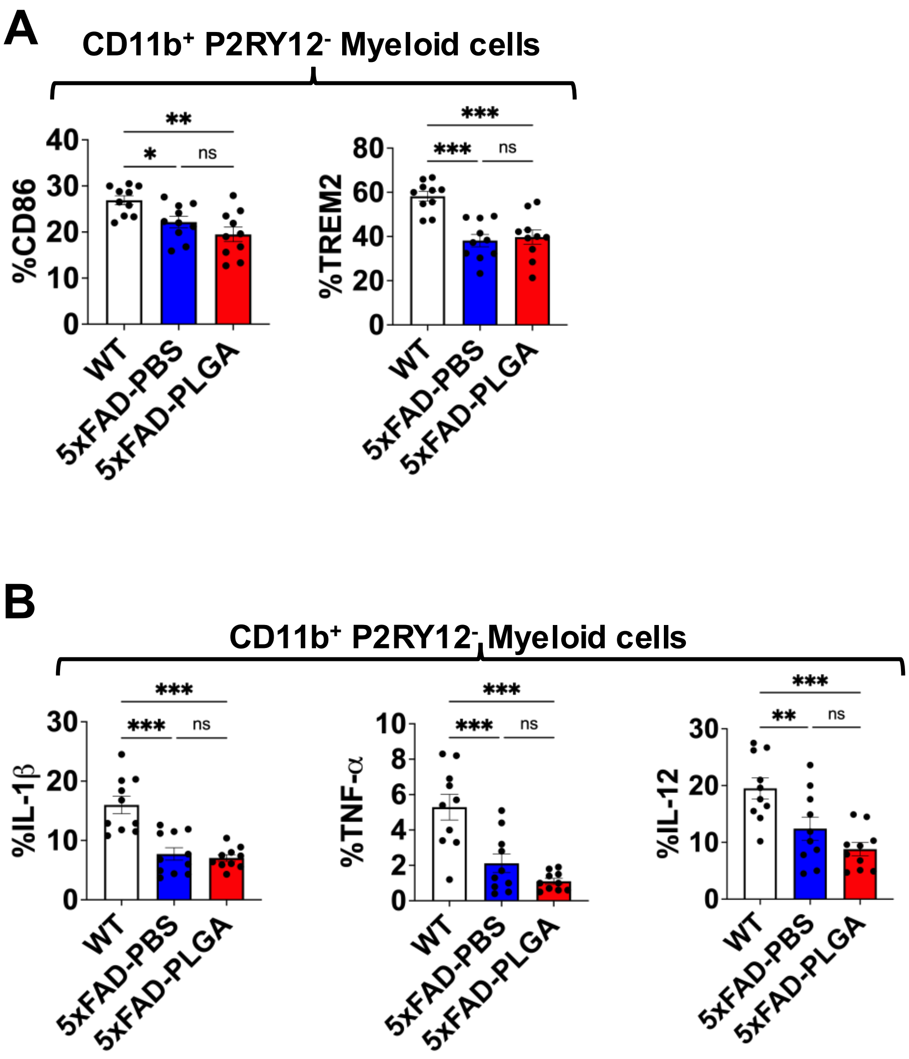
A
CD11b+ P2RY12- Myeloid cells
%CD86
%TREM2
**
*
ns
***
***
ns
WT
5xFAD-PBS
5xFAD-PLGA
B
CD11b+ P2RY12- Myeloid cells
%IL-1β
%TNF-α
%IL-12
***
***
ns
***
***
ns
***
**
ns
WT
5xFAD-PBS
5xFAD-PLGA

Figure S4

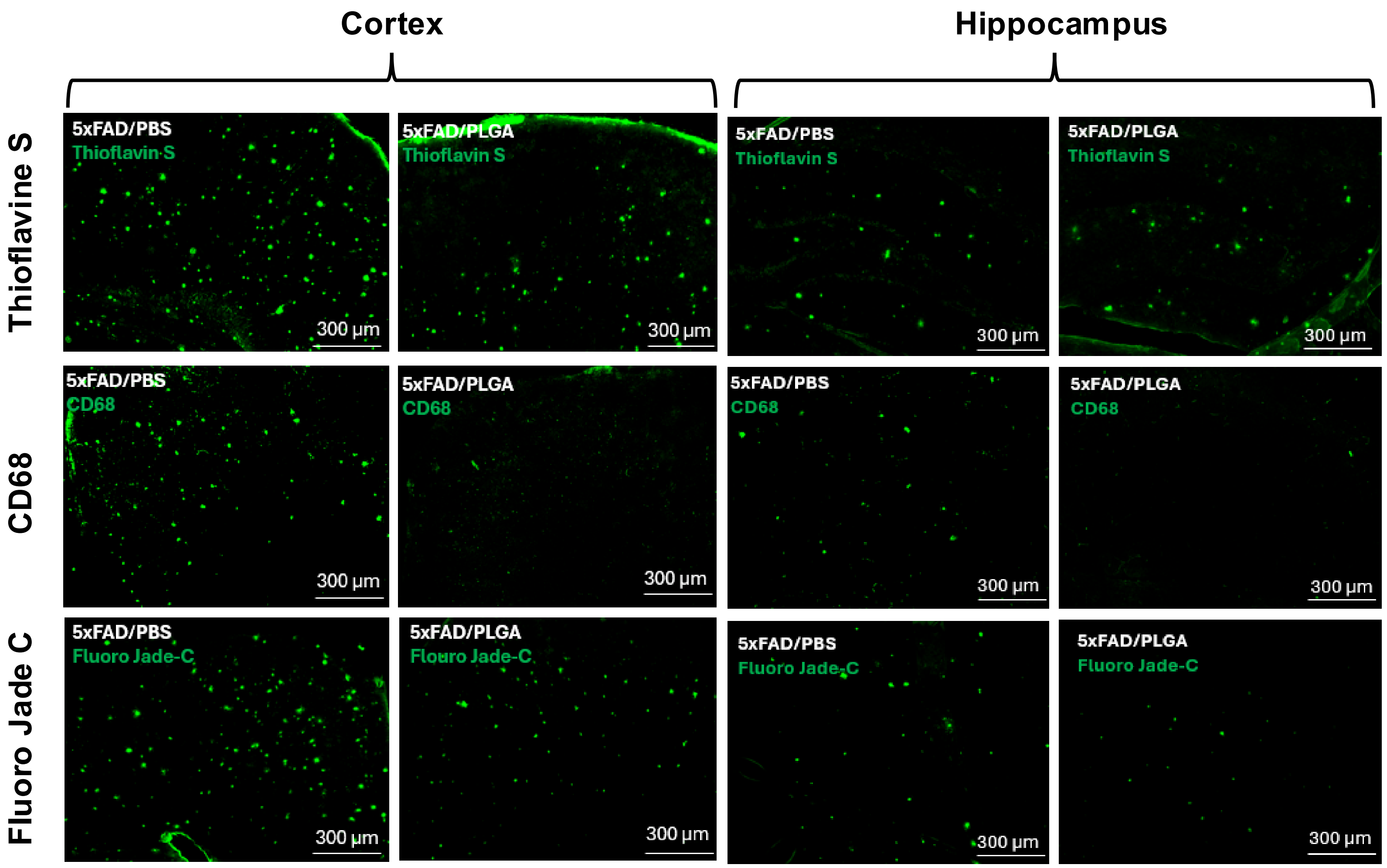
Cortex
Hippocampus
Thioflavine S
CD68
Fluoro Jade C
5xFAD/PBS
Thioflavin S
5xFAD/PLGA
Thioflavin S
5xFAD/PBS
CD68
5xFAD/PLGA
CD68
5xFAD/PBS
Fluoro Jade-C
5xFAD/PLGA
Flouro Jade-C
5xFAD/PLGA
Fluoro Jade-C
300 μm
